# Pyrophosphate modulates stress responses via SUMOylation

**DOI:** 10.1101/504373

**Authors:** M. Görkem Patir-Nebioglu, Zaida Andrés, Melanie Krebs, Fabian Fink, Katarzyna Drzewicka, Nicolas Stankovic-Valentin, Shoji Segami, Sebastian Schuck, Michael Büttner, Rüdiger Hell, Masayoshi Maeshima, Frauke Melchior, Karin Schumacher

## Abstract

Pyrophosphate (PPi), a byproduct of macromolecule biosynthesis is maintained at low levels by soluble inorganic pyrophosphatases (sPPase) found in all eukaryotes. In plants, H+-pumping pyrophosphatases (H+-PPase) convert the substantial energy present in PPi into an electrochemical gradient. We show here, that both cold- and heat stress sensitivity of *fugu5* mutants lacking the major H+-PPase isoform AVP1 is caused by reduced SUMOylation. In addition, we show that increased PPi concentrations interfere with SUMOylation in yeast and we provide evidence that SUMO activating E1-enzymes are inhibited by micromolar concentrations of PPi in a non-competitive manner. Taken together, our results do not only provide a mechanistic explanation for the beneficial effects of AVP1 overexpression in plants but they also highlight PPi as an important integrator of metabolism and stress tolerance.

## INTRODUCTION

Reshaping of metabolic networks under stress conditions enables the synthesis of protective compounds while metabolic homeostasis needs to be maintained. In about 200 metabolic reactions ATP is not used as a phosphorylating but as an adenylating reagent leading to the release of inorganic pyrophosphate (PPi). Most prominently, the biosynthesis of many macromolecules including DNA, RNA, proteins and polysaccharides releases large amounts of PPi (Ferjani et al., 2014; Heinonen, 2001). Given the substantial free energy of PPi, the efficient biosynthesis of macromolecules requires that PPi is immediately destroyed to prevent the respective back-reactions (Kornberg, 1962). In all eukaryotes PPi is hydrolysed by soluble inorganic pyrophosphatases (sPPase; EC 3.6.1.1) in a highly exergonic reaction. Loss of sPPase function causes lethality in yeast (Lundin et al., 1991) and C. elegans (Ko et al., 2007) presumably due to accumulation of PPi inhibiting the biosynthesis of macromolecules. Arabidopsis encodes six sPPase-paralogs (PPa1-PPa6) of which only PPa6 is localized in plastids whereas all others are cytosolic (Gutiérrez-Luna et al., 2016; Segami et al., 2018). However, their PPase activity is rather low and even the loss of the four ubiquitously expressed isoforms does not cause phenotypic alterations (Segami et al., 2018). In contrast, expression of E. coli sPPase severely affects plant growth via alterations in carbon partitioning between source and sink organs caused by the inhibition of several plant enzymes involved in carbohydrate metabolism that use PPi as an energy source (Geigenberger et al., 1998; Sonnewald, 1992). Importantly, in addition to soluble PPases, plants contain membrane-bound proton-pumping pyrophosphatases (H^+^-PPase) at the tonoplast and in the Golgi that convert the energy otherwise lost as heat into a proton-gradient (Maeshima, 2000; Segami et al., 2010). Fugu5 mutants lacking the tonoplast H^+^-PPase AVP1 were identified based on their phenotype characterised by compensatory cell enlargement due to a decrease in cell number (Ferjani et al., 2011). The fact that the *fugu5* phenotype could be rescued either by growth in the presence of exogenous sucrose or the expression of the yeast sPPase IPP1 showed clearly that altered PPi levels and not reduced H+-pumping are causative (Asaoka et al., 2016; Ferjani et al., 2011). Indeed, vacuolar pH is only mildly affected in *fugu5* mutants indicating that the H^+^-pumping ATPase (V-ATPase) present at the tonoplast is largely sufficient for vacuolar acidification (Ferjani et al., 2011; Kriegel et al., 2015). However, loss of both vacuolar proton-pumps leads to a much more severe phenotype and defect in vacuolar acidification than loss of the tonoplast V-ATPase alone (Kriegel et al., 2015). It has indeed been discussed that AVP1 serves as a backup system for the V-ATPase in particular under ATP-limiting conditions like anoxia or cold stress (Maeshima, 2000). During cold acclimation plants accumulate cryoprotectants including sugars in their vacuoles and activity of both proton-pumps is upregulated leading to improved freezing tolerance (Schulze et al., 2012; Thomashow, 1999). Overexpression of AVP1 has been shown to cause increased plant growth under various abiotic stress conditions including salinity, drought and phosphate starvation but the underlying mechanism remained unclear (Gaxiola et al., 2012; Park et al., 2005; Schilling et al., 2017). Attachment of the small ubiquitin-related modifier SUMO to substrate proteins plays a central role in the response to a broad set of stress responses including the ones affected by AVP1 overexpression (Castro et al., 2012). Modification of target proteins by SUMO-conjugation proceeds via a three-step mechanism. First the SUMO moiety is adenylated and then bound via a high-energy thioester linkage to the heterodimeric SUMO-activating enzyme (E1) leading to the release of PPi. Next, the activated SUMO is transferred to the SUMO-conjugating enzyme E2 and finally, assisted by SUMO-protein ligase (E3), donated to a large set of substrate proteins (Flotho and Melchior, 2013; Johnson, 2004). In Arabidopsis, the key transcriptional regulator of the cold response INDUCER OF CBF EXPRESSION 1 (ICE1) as well as the heat shock factor A2 (HSFA2) have been shown to be positively regulated by SUMOylation (Cohen-Peer et al., 2010; Miura et al., 2007). In this study, we report that AVP1 contributes to both cold acclimation and heat tolerance and we show that the rapid increase in SUMOylation common to both stress responses is missing in the absence of AVP1. Furthermore, we provide evidence that accumulation of PPi in plants, yeast and mammals inhibits the SUMO E1 activating enzyme in turn affecting the fate, localization or function of a large number of proteins during cellular stress responses. Our results provide a mechanistic explanation for the beneficial effects of AVP1 overexpression in plants and highlight PPi as an important integrator of metabolism and stress tolerance.

## RESULTS

### Lack of V-PPase activity impairs cold acclimation

We have shown previously that upregulation of ATP-hydrolysis by the V-ATPase during cold acclimation depends on the presence and the activity of the V-PPase (Kriegel et al., 2015). To complete the data-set for vacuolar proton-pump activity during cold acclimation, we performed parallel measurements of ATP- and PPi-hydrolysis, H^+^-pumping as well as cell sap pH in wild-type (Col-0), the *fugu5–1* mutant and a UBQ:AVP1 overexpression line. Both ATP- and PPi-dependent proton-pumping are increased in wt and UBQ:AVP1 during cold-acclimation (Supplemental Figure 1A+B). As expected PPi-dependent proton-pumping was undetectable in *fugu5–1*, but ATP-dependent proton-pumping was also reduced in *fugu5–1* compared to wt and increased only marginally upon cold-acclimation (Supplemental Figure 1B). As a consequence of cold-induced proton-pump stimulation, vacuolar pH drops by 0.1 pH-units in wt and UBQ:AVP1 but not in *fugu5–1* (Supplemental Figure 1C).

**Figure 1.**
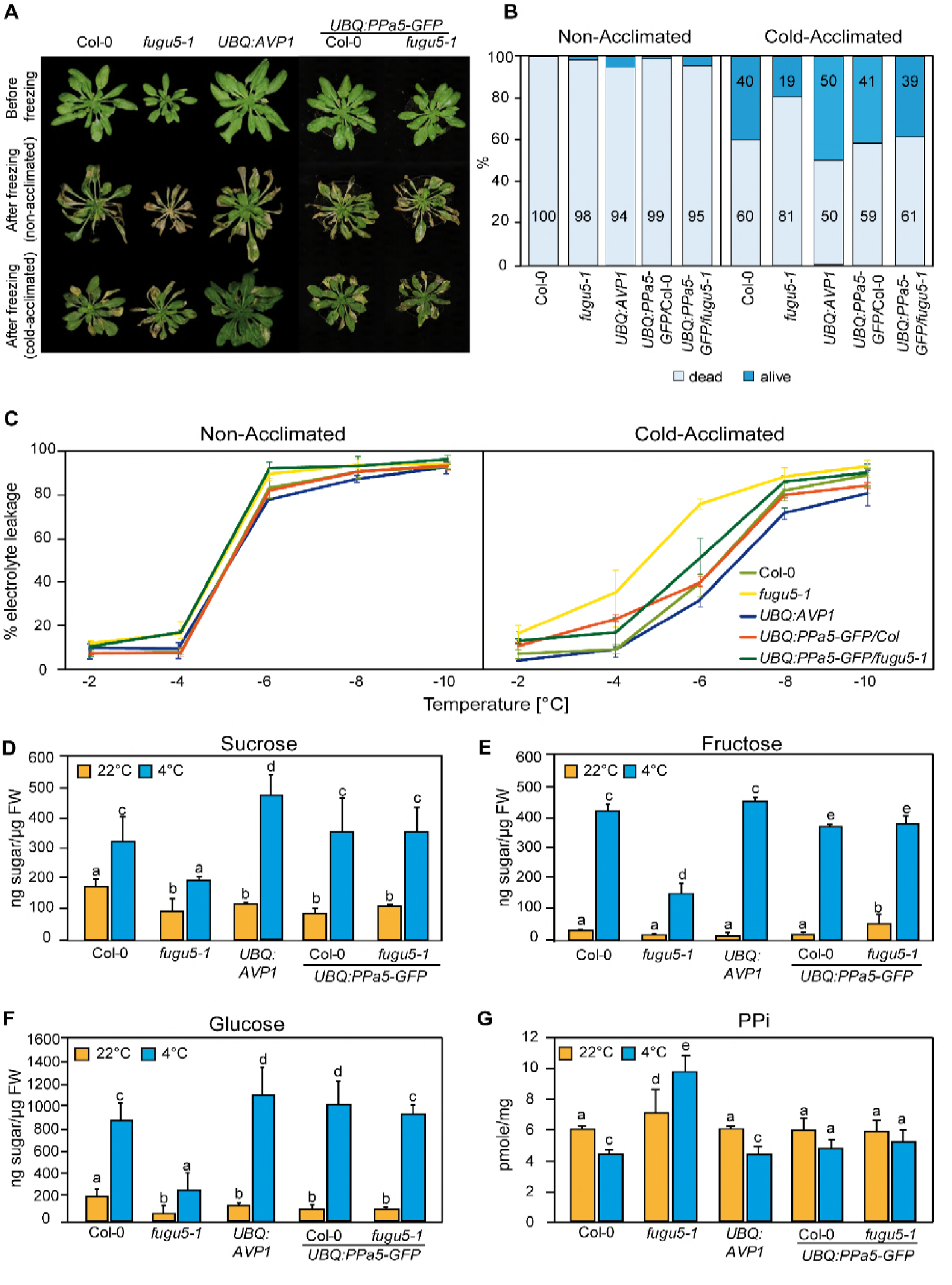
PP_i_ hydrolysis is required to rescue the freezing sensitive phenotype of *fugu5–1*. **(A)** and **(B)** Freezing tolerance assay. Wt, *fugu5–1, UBQ:AVP1* and *UBQ:PPa5-GFP* in wt and *fugu5–1* background were grown for 6 weeks at 22 °C and were then moved to 4°C for cold-acclimation, or kept at 22 °C for 4 days. Afterwards plants were subjected to a 5-h freezing temperature regime (0 to −10 °C). After thawing at 4 °C overnight, plants were moved back to 22 °C. **(A)** Images were taken before cold-acclimation and one week after the freezing treatment. **(B)** Quantification of dead and alive leaves was done one week after the freezing treatment with n ≥ 75 leaves. 3 independent experiments were performed. **(C)** Electrolyte leakage assay of Wt, *fugu5–1, UBQ:AVP1* and *UBQ:PPa5-GFP* in Wt and *fugu5-1* background was performed on leaf material of acclimated and non-acclimated plants at indicated freezing temperatures. Error bars represent SD of the mean of n=3 biological replicates. **(D-G)** Sugar and PPi measurements were done from extracts of acclimated (4 °C) and non-acclimated (22 °C) 6-week old rosette leaves. Error bars show SD of the mean with n=3 samples of one representative experiments. 3 biological replicates were performed. Significant differences are indicated by different letters (Two-way ANOVA followed by Tukey’s test, p<0.05).

Whereas the seedling phenotype of *fugu5–1* is rescued by expression of the yeast soluble PPase IPP1 under control of the AVP1-promoter during the seedling stage (Ferjani et al., 2011), the adult growth phenotype of plants grown in short day (SD) was not rescued (Supplemental Figure 2A). We thus expressed the constitutively expressed Arabidopsis soluble pyrophosphatase PPa5 fused to GFP under the control of the UBQ10-promoter and could show that it is located in the cytosol as well as in the nucleus (Supplemental Figure 2B) and fully rescues the seedling (Supplemental Figure 2C) as well as the adult phenotype (Supplemental Figure 2D+E) of *fugu5–1*. For further analysis, two lines in the wild-type and in the *fugu5–1* background that showed protein expression of PPa5-GFP (Supplemental Figure 2F) and comparable increased total soluble pyrophosphatase activity (Supplemental Figure 2G) in the wild-type and in the *fugu5–1* background were chosen. We next asked if cold-acclimation is affected in *fugu5–1* and if so, whether this could be rescued by overexpression of a soluble pyrophosphatase.

**Figure 2.**
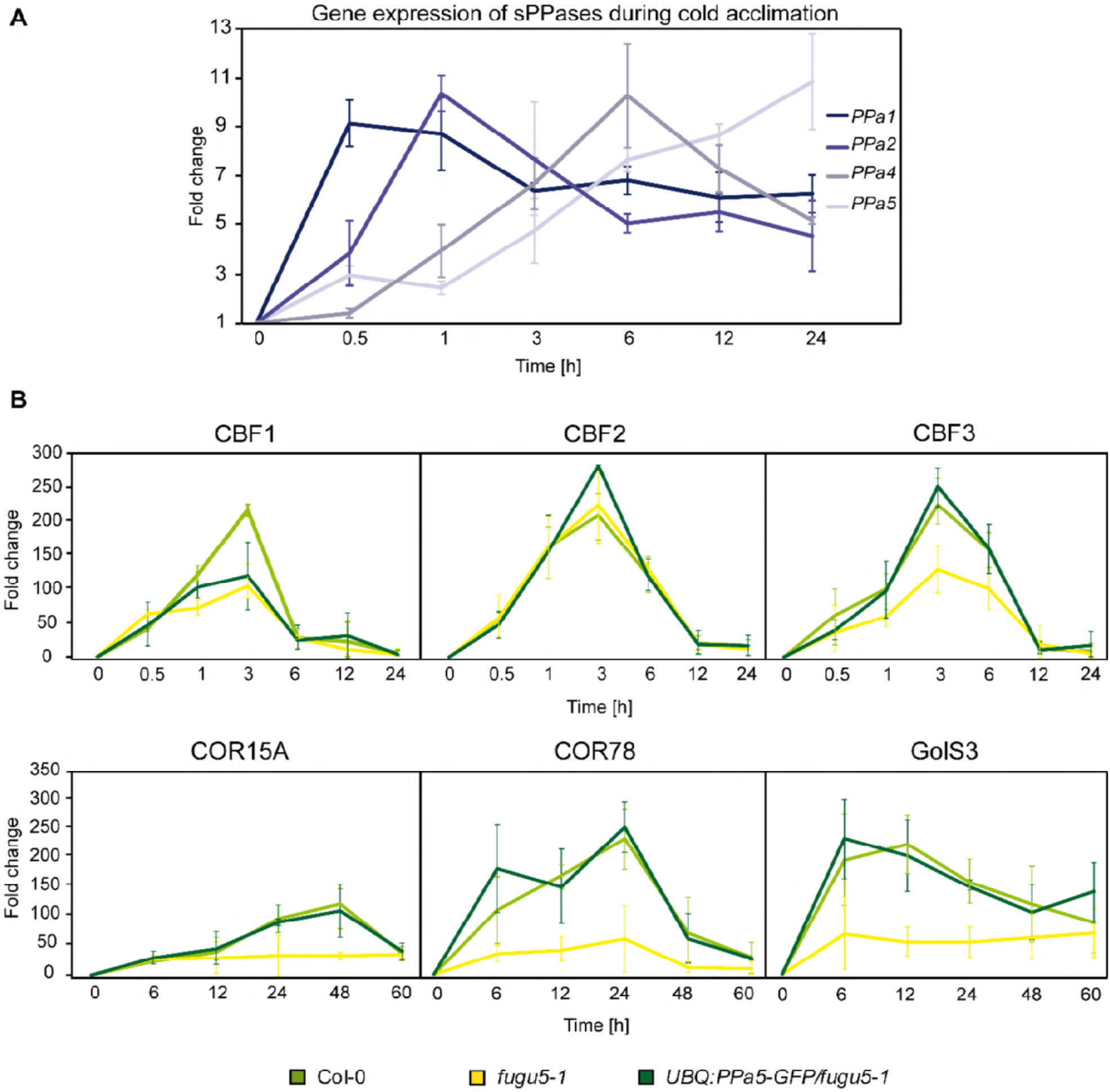
sPPase expression is induced upon cold exposure to control PP_i_ that affects expression of CBFs and CBF target genes. **(A)** qRT-PCR for the analysis of expression of sPPase 1,2,4 and 5. **(B)** Measurement of expression of *CBF, COR* and *GolS3* genes in Col-0, *fugu5–1* and *UBQ:PPa5-GFP/fugu5–1* by qRT-PCR. **(A)** and **(B)** Plants were grown for six weeks under short-day conditions at 22°C. Afterwards, they were exposed to 4°C for indicated time periods. Whole rosettes were used for total RNA extraction. *Actin2* expression was used for normalization. Error bars represent SD of the mean of n=3 biological replicates. Data analysis was performed using the ΔΔC_t_ method.

Both survival rate and ion release as a measure of freezing tolerance was comparable in all genotypes exposed to freezing without prior cold acclimation (Figure 1A + B). Cold acclimation via exposure to 4°C for 4 days significantly improved freezing tolerance in wild-type to a much higher extent than in *fugu5–1* plants and expression of PPa5 fully rescued the hypersensitivity to cold (Figure 1C). Accumulation of soluble sugars during exposure to low temperatures contributes to freezing tolerance and could be directly affected by PPi-accumulation (Ferjani et al., 2018). We thus next compared the accumulation of glucose, fructose and sucrose and found that cold-induced sugar accumulation is indeed strongly reduced in the *fugu5–1* mutant but restored by UBQ:PPa5-GFP (Figure 1D-1F) suggesting that accumulation of PPi and not a lack of H^+^-pumping is responsible for the impaired cold acclimation in *fugu5–1*. In agreement with this hypothesis, we found that PPi levels are reduced in the wt during cold-acclimation whereas they increase in *fugu5–1* resulting in 2-fold higher levels compared to wt after cold acclimation (Figure 1G).

### PPi controls cold-acclimation via SUMOylation

Low temperature triggers the expression of the CBF (C-repeat binding factor) family of transcription factors, which in turn activate downstream genes that confer chilling and freezing tolerance (Chinnusamy et al., 2007). We used qRT-PCR to profile the expression of members of the PPa-gene family over 24h after exposure to low temperature (4°C) and found that *PPa1* and *PPa4* are rapidly induced after cold exposure whereas transcripts of *PPa2* and *PPa5* accumulated at later time points (Figure 2A). Upregulation of sPPase genes suggests that PPi-levels are actively controlled during the early cold acclimation response and we thus next compared the expression levels of the core transcriptional regulators *CBF1–3* as well as the three downstream response genes COR15A, COR78 and GolS3. Whereas expression of *CBF2* was nearly unaffected, *CBF1* and in particular *CBF3* induction was found to be strongly reduced in the *fugu5–1* mutant (Figure 2 B). Similarly, induction of all three target genes was found to be strongly reduced throughout the cold response (Figure 2 C). Cold-induction was restored when PPa5 was constitutively expressed in the *fugu5–1* background (Figure 2 B+C) indicating that the observed changes in gene expression are caused by reduced PPi-hydrolysis and not by reduced H^+^-pumping.

The fast transcriptional response to cold is initiated by ICE1 (Inducer of CBF expression 1), a direct activator that is negatively regulated by ubiquitination-mediated proteolysis and positively regulated by SUMOylation (Dong et al., 2006; Miura et al., 2007); Figure 3 A). Using a specific antibody to detect ICE1 in total seedling protein extracted in the presence of NEM to inhibit deSUMOylation, we observed two bands in wild-type that are both absent in *ice1–2* indicating that they correspond to a non-modified (100kD) and modified (130kD) dimer of ICE1. The ICE1 monomer (50kD) was only observed when proteins were extracted in the presence of DTT and without NEM (Supplemental Figure 3).

**Figure 3.**
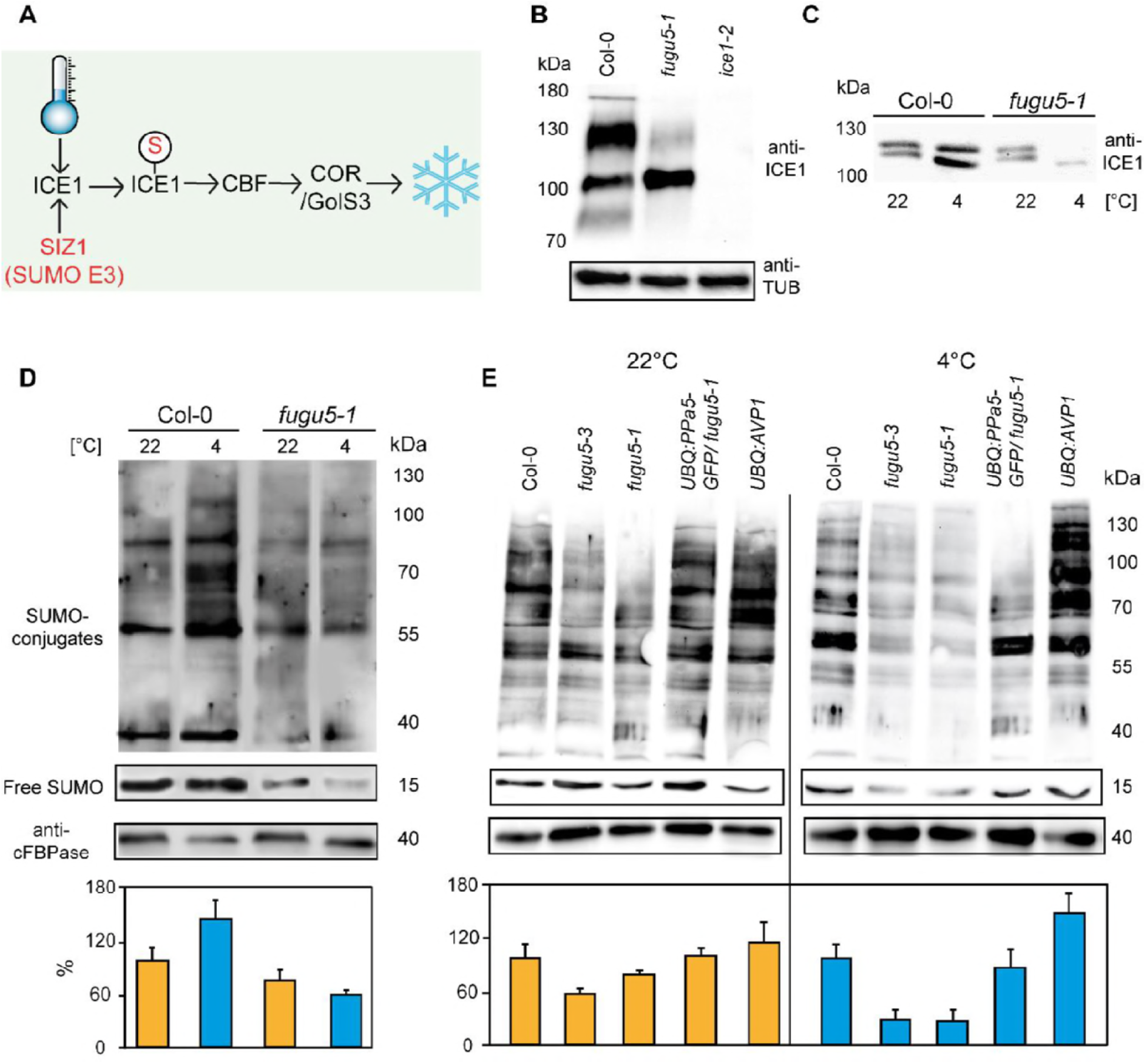
Modification of ICE1 and general SUMOylation upon cold exposure are inhibited *fugu5*. **(A)** Cold acclimation induces ICE1 sumoylation which then activates CBFs and leads to the expression downstream targets for the establishment of freezing tolerance. **(B)** Determination of the amount of ICE1 in Col-0, *fugu5–1* and *ice1–2* seedlings. Amount of TUBULIN was detected as loading control. (C) Comparison of the amount of the ICE1 under normal conditions (22°C) and after cold treatment (4°C, 3h) in Col-0 and *fugu5–1* seedlings. **(B)** and **(C)** 10 days old liquid grown seedlings were used for total protein extraction. Anti-ICE1 was used as primary antibody. **(D)** Western blots comparing sumoylation levels of Col-0 and *fugu5–1* under normal conditions (22°C) and after cold treatment (4°C, 3h). **(E)** Western blots demonstrating the total sumoylation in Wt, *fugu5–1, fugu5–3, UBQ:AVP1* and *UBQ:PPa5-GFP* under normal conditions (22°C) and after cold treatment (4°C, 3h). **(D)** and **(E)** 10 days old liquid grown seedlings were used for total protein extraction. Anti-SUMO1/2 was used as primary antibody. Whole lanes were measured for the calculation of protein amounts using ImageJ. cFBPase detection was used for normalization. Error bars represent SD of n≥2 biological replicates.

In *fugu5–1* the modified dimer was barely detectable indicating that either ubiquitination or sumoylation of ICE1 are affected (Figure 3 B). We next compared levels of ICE1 during cold acclimation and found that ICE1 accumulated after exposure to 4°C for 3h in wild-type but was strongly reduced in *fugu5–1* (Figure 3 C). We thus next asked if overall cold-induced SUMOylation was affected in *fugu5*. Whereas cold exposure let to a rapid and massive accumulation SUMO1/2 conjugates in the wild-type, this response was absent in *fugu5–1* (Figure 3 D). Quantification of SUMO levels in two independent fugu5-alleles showed that it is already reduced to 60% of wt in plants grown at 22°C and levels drop to 20% of wt after incubation at 4°C for 3h (Figure 3 E). SUMOylation is restored to wild-type levels in UBQ:PPa5-GFP and slightly enhanced in UBQ:AVP1 plants indicating that low PPi levels are critical for efficient SUMOylation (Figure 3 D).

### PPi inhibits heat-stress induced SUMOylation in both plants and yeast

Rapid and reversible accumulation of SUMO conjugates does not only occur during cold stress but also during heat stress (Miller et al., 2010; Rytz et al., 2018) and accumulation of PPi should thus also inhibit the heat stress response. Indeed, survival rate of *fugu5* seedlings was strongly reduced by exposure to 40°C for 30 min but restored in the UBQ:PPa5 complementation line. Of note, the survival rate of the UBQ:AVP1 overexpression line was increased compared to the wt (Figure 4 A and 4B). We therefore analysed next if heat induced SUMO1/2 conjugate accumulation was affected. Exposure to 40°C for 30 min led to accumulation of SUMO1/2 conjugates in the wt, whereas the response was strongly reduced in *fugu5–1* (Figure 4 C). Consistently, a reduction of SUMO levels after heat stress was also observed for fugu5–3 are reduced to 20% of wt after incubation 40°C for 30’ in both *fugu5–1* and *fugu5–3* but was restored to wt levels in UBQ:PPa5-GFP and UBQ:AVP1 plants (Figure 4 D).

**Figure 4.**
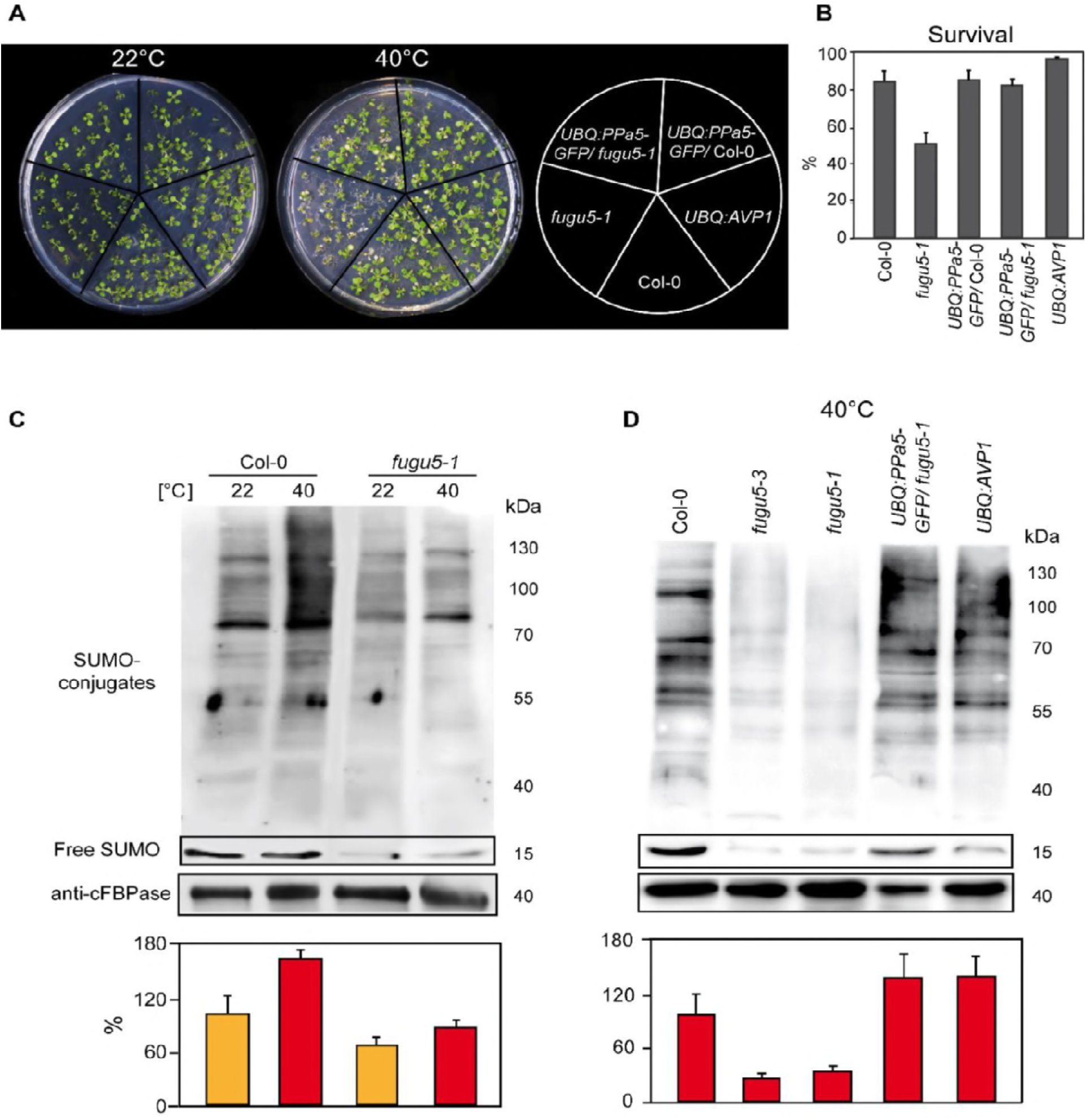
Heat shock-induced SUMOylation is also reduced in *fugu5*. **(A)** Phenotypic analysis of 10 days old Col-0, *fugu5–1, UBQ:PPa5-GFP* in Col-0 and *fugu5–1* backgrounds and *UBQ:AVP1* seedlings analysis before and after heat. Representative pictures of seedlings before and 4 days after completion of heat shock treatment are depicted. **(B)** Seedling survival was determined 4 days after the heat shock. Alive and dead seedlings were counted and survival is shown as the percentage of the living seedlings. Error bars show SD of the mean with n≥24 samples of one representative experiments. 2 biological experiments were performed. **(C)** Sumoylation levels of Col-0 and V-PPase mutant *fugu5–1* were analysed with western blot under normal conditions (22°C) and after heat shock treatment (40°C, 30 min). **(D)** Measurement of the SUMO amount of Col-0, V-PPase mutants, *UBQ:PPa5-GFP/fugu5–1* and *UBQ:AVP1* seedlings after heat shock treatment (40°C, 30 min). **(C)** and **(D)** 10 days old liquid grown seedlings are used for total protein extraction. Anti-SUMO1/2 (Agrisera) was used as primary antibody. Whole lanes were measured for the calculation of protein amounts using ImageJ. cFBPase detection was used for normalization. Error bars represent SD of n=2 biological replicates.

SUMO plays an important role in stress responses across all eukaryotes (Enserink, 2015; Hannich et al., 2005). Therefore, we asked whether PPi accumulation has a comparable effect in the yeast S. *cerevisiae*. We employed a strain in which the sole and essential sPPase IPP1 is expressed under the control of the *GAL1* promoter (Serrano-Bueno et al., 2013) so that switching the carbon source from galactose to glucose led to a depletion of IPP1 (Figure 5 A) after 6h that was almost complete after 15h (Figure 5 A). When wt yeast was subjected to heat stress (40 °C, 1 h), SUMOylation increased by a factor of two (Figure 5B). Heat-induced SUMOylation was strongly diminished by depletion of IPP1 depletion phase (Figure 5C) indicating that inhibition of SUMOylation by PPi is not limited to plants.

**Figure 5.**
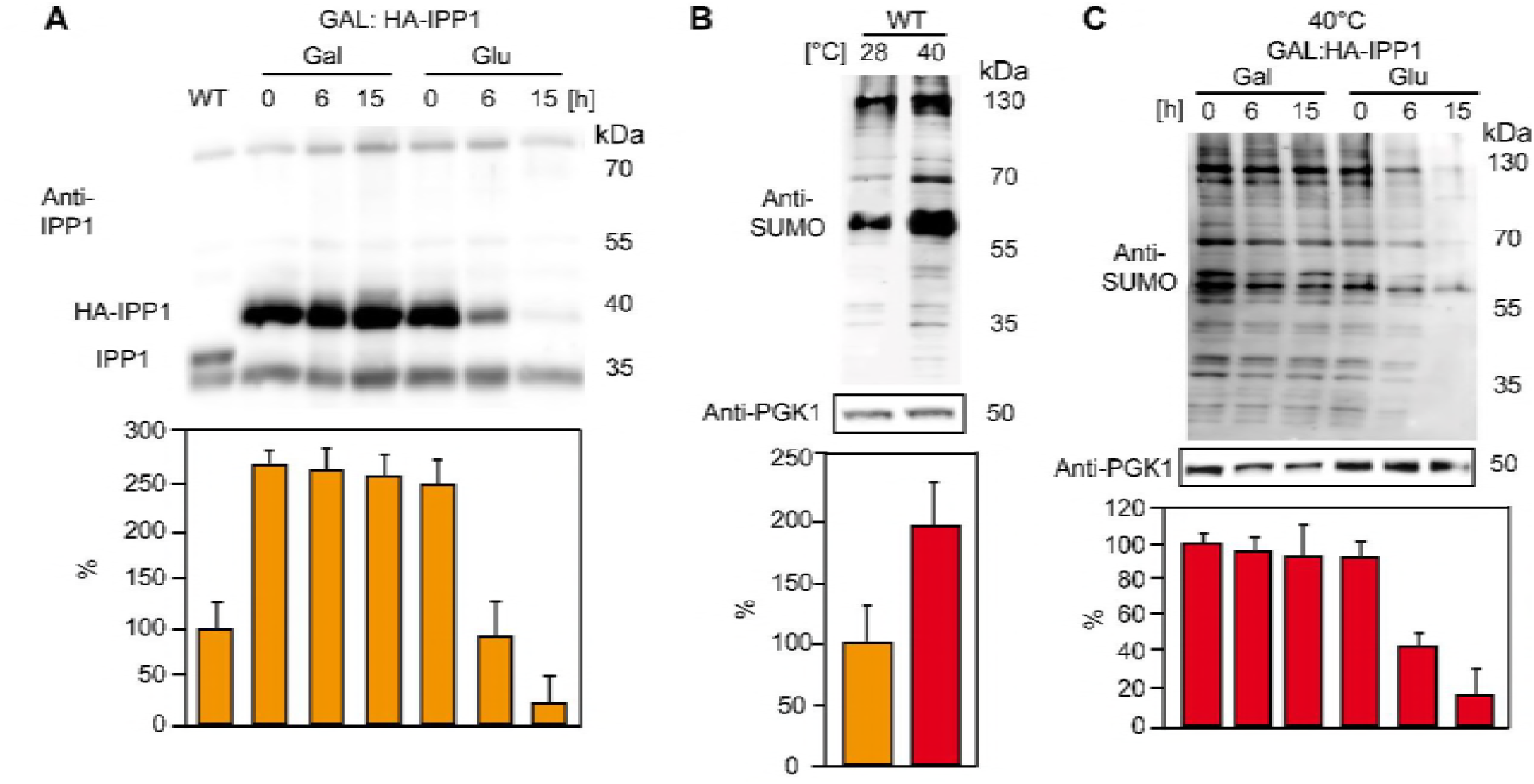
Increased PPi levels interfere with SUMOylation in yeast. **(A)** Amount of the soluble pyrophosphatase protein in a conditional Ipp1 mutant of S. *cerevisiae* (GAL:HA-IPP1). Wt strain (W303) is used as a control. **(B)** Amount of total sumoylation in W303 determined before and after heat stress. **(C)** Measurement of total SUMO protein in conditional IPP1 mutant of S. *cerevisiae* (GAL:HA-IPP1). **(A-C)** Yeast is grown in synthetic complete medium supplemented with galactose at 28°C. After growing until OD_600_ 0.5, part of the Ipp1 conditional mutant is switched to glucose supplemented medium to suppress the promoter and samples are collected at the indicated time points. For the heat treatment, cultures are switched to 40°C incubator for 1 hour. Error bars represent SD of n≥2 biological replicates.

### What is the mechanistic link between PPi accumulation and SUMOylation?

Conjugation of SUMO to target proteins is initiated by E1 enzymes through adenylation, a reaction that releases PPi and could thus be inhibited by increased cytosolic PPi levels (Haas et al., 1982; Lois and Lima, 2005). To test the direct effect of PPi on SUMOylation, we employed an in vitro assay in which conjugation of YFP-SUMO to RanGAP1-CFP can be measured as a change in FRET efficiency if the E1 and E2 enzymes as well as ATP are provided (Bossis et al., 2005). Addition of micromolar concentrations of PPi caused a strong inhibition of RanGAP1-CFP SUMOylation by the human E1 (Uba2/Aos1) and E2 (Ubc 9) which could be released by addition of a soluble pyrophosphatase (Figure 6B + 6C). To determine the mode of inhibition, we determined Vmax and Km in the absence as well as in the presence of PPi (Figure 6B) leading to the conclusion that inhibition of E1E2 activity by PPi follows a mixed mode (Figure 6D). We purified the Arabidopsis E1 heterodimer SAE1b SAE2 but could not detect activity in the FRET assay (Supplemental Figure 3) and thus analysed SAE2~SUMO thioester formation in the presence and absence of PPi. In accordance with the results for the human enzyme, the Arabidopsis SUMO E1-activity is inhibited by 10uM PPi (Figure 6E).

**Figure 6.**
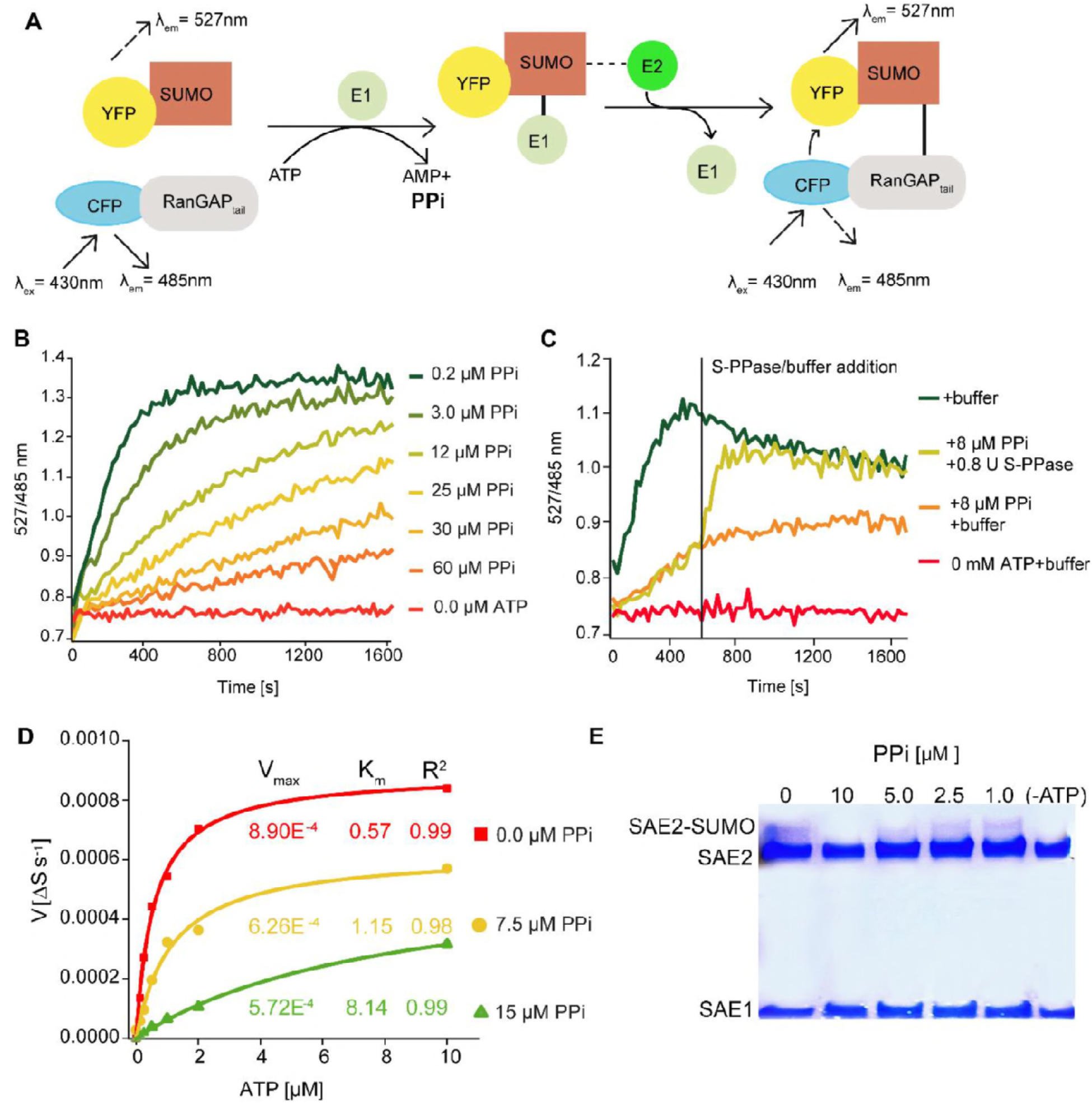
PPi regulates SUMOylation activity *in vitro*. **(A)** Schematic illustration of the FRET-based sumoylation assays. Upon addition of ATP, the human SUMO E1 activating enzyme Aos1/Uba2 and the E2 conjugating enzyme Ubc9 form an isopeptide bond between the CFP-tagged human model substrate GAP_tail_ and YFP-tagged mature SUMO2. This can be detected via FRET measurements: Following the excitation of CFP, energy is transferred onto YFP. YFP and CFP emission are recorded upon excitation at 430 nm. Measurements are calculated as the ratio of the λ_em_ (SUMO2-YFP, 527 nm) to λ_ex_ (CFP-GAP_tail_, 485 nm). (B) PP_i_ titration showing that the increasing PP_i_ concentration inhibits the sumoylation activity. 1mM ATP used for all the measurements. **(C)** *In vitro* sumoylation assay showing that the *E. coli* soluble PPase is able to remove the inhibitory effect of PPi. After 10 min of measurement, one of the 8 μM PPi containing wells were supplied with 0.8 U of *E.coli* soluble PPase and control buffer was added to the rest of the wells. Measurements were continued for 20 more minutes. **(D)** PP_i_ addition results in mixed inhibition of sumoylation activity. Michaelis-Menten fittings of the measurements shown in Supplement figure 3. Fittings are done in Origin software and V_max_ and K_m_ are calculated accordingly. **(E)** *In vitro* thioester bond formation assay showing Arabidopsis E1 (SAE2/SAE1a) and SUMO conjugation under different PPi concentrations. **(B-D)** Experiments were repeated 4 times, one representative image is shown.

## DISCUSSION

It has long been assumed that the combined action of V-ATPase and V-PPase enables plants to maintain transport into the vacuole even under stressful conditions (Maeshima, 2000). We have shown previously that the increased activity of the V-ATPase during cold acclimation is largely dependent on the presence of the V-PPase (Kriegel et al., 2015). During cold acclimation *fugu5* mutants thus should not able to adjust their tonoplast proton-pumping activity to the increased demand caused by the accumulation of soluble sugars, organic acids and other osmoprotectants in the vacuole (Schulze et al., 2012). We show here that lack of the V-PPase indeed limits cold acclimation severely. However complementation by overexpression of the soluble pyrophosphatase PPa5 shows clearly that this phenotype is not caused by a reduced proton-gradient limiting cold-induced accumulation of solutes in the vacuole (Figure 1). Accumulation of PPi has been shown to be causative for the developmental phenotype of fugu5 seedlings (Ferjani et al., 2018, 2011) and our results show that this also applies to the freezing tolerance and heat stress phenotypes caused by the lack of AVP1 that we report here for the first time. Although overexpression of AVP1 has been shown to result in increased stress tolerance and yield in multiple crop plants, reduced stress tolerance of *fugu5* mutants has so far not been reported. The fact that the seedling phenotype observable during the heterotrophic phase of *fugu5* seedlings could be rescued by supply of exogenous sucrose pointed to an inhibition of gluconeogenesis. The Glc1P/UDP-Glc reaction is reversible and it has been shown that UGP-Glc pyrophosphorylase is a major target of PPi-inhibition during seedling establishment (Ferjani et al., 2018). Similarly, PPi accumulation could inhibit sugar accumulation during cold acclimation but the fact that the early transcriptional response to cold is dampened in the *fugu5* mutant is not easily explained solely by a shift in sugar metabolism (Gutiérrez-Luna et al., 2018). PPi is not only released by many anabolic reactions but also by E1 enzymes that initiate the attachment of ubiquitin or ubiquitin-like proteins (UBLs) including SUMO. Activation of UBLs requires ATP and occurs via carboxy-terminal adenylation and thiol transfer leading to the release of AMP and PPi and would thus be prone to inhibition by PPi accumulation (Desterro et al., 1999; Schulman and Harper, 2009). The MYC-like bHLH transcriptional activator ICE1 is subject to ubiquitination-mediated proteolysis under ambient temperature that is counteracted by SUMOylation during the cold response (Miura and Hasegawa, 2008). We have shown here that the compromised cold acclimation of *fugu5* is caused by the failure to stabilize ICE1 and that the overall levels of SUMO-conjugates that rapidly increase upon cold exposure in the wild-type fail to increase in *fugu5* (Figure 3). As we cannot exclude that the altered sugar metabolism of fugu5 indirectly impinges SUMOylation during cold acclimation, we extended our analysis to the heat stress response. The rapid and reversible accumulation of SUMO conjugates is one of the fastest molecular responses observed during heat stress (Kurepa et al., 2003; Rytz et al., 2018). The fact that this response is dampened in both plants and yeast when PPi accumulates (Figures 4 and 5) argues strongly against a secondary metabolic effect. Evidence for a direct inhibitory effect of PPi on SUMOylation was obtained in an *in vitro* FRET-based assay that allowed us to determine that the SUMOylation of RanGAP catalysed by human E1 and E2 enzymes was inhibited by micromolar concentrations of PPi following a mixed mode of inhibition (Figure 6). Although we cannot exclude that PPi could inhibit the action of the E2 enzyme, the reaction catalysed by the heterodimeric E1 activating enzyme releases PPi and is thus most likely inhibited when PPi accumulates. Indeed, we could show that E1 subunit SAE2~SUMO thioester formation is inhibited in the presence of micromolar PPi, raising the question how exactly PPi inhibits E1-activity (Figure 6).

For adenylation of the SUMO C-terminus to occur, the E1 enzyme adopts an open conformation that allows binding of ATP. In this conformation, the catalytic cysteine of E1 is too far away and unavailable to become linked to SUMO. Thioester bond formation between E1 and SUMO requires structural remodelling to a closed conformation in which the catalytic cysteine moves adjacent to the C terminus of SUMO~AMP, via unfolding of structures associated with ATP binding and SUMO adenylation (Lois and Lima, 2005; Olsen et al., 2010). It has been suggested that active site remodelling pushes the E1 reaction forward by promoting the release of pyrophosphate to prevent the reverse reaction, the attack of the adenylate by pyrophosphate leading to the reformation of ATP. Not only is the adenylation step rate limiting, once the thioester bond is formed and AMP is released, E1 switches back to the open conformation and a second adenylation reaction occurs, resulting in the formation of a ternary complex, with an E1 molecule binding to one SUMO molecule at the adenylation active site and to a second via a thioester bond through the catalytic cysteine (Olsen et al., 2010). As E1 enzymes are potential targets for therapeutic intervention in cancer and other diseases understanding their enzymatic activity as well as inhibitory mechanisms at the atomic level may provide leads for the development of novel drugs. A novel allosteric inhibitor that targets a cryptic pocket distinct from the active site and locks the enzyme in a previously unobserved inactive conformation has recently been identified (Lv et al., 2018) and it will be of great interest to determine how accumulation off PPi affects the conformation of E1.

Although the exact mechanism remains to be determined, the fact that E1 activity is classically measured as ATP:PPi (Haas et al., 1982; Haas and Rose, 1982) exchange clearly reflects that inhibition of E1 enzymes by PPi is not novel per se. Although cytosolic PPi concentrations reported in the literature, in particular for plants, strongly suggest that relevant concentrations occur not only in mutant backgrounds or under stress conditions the relevance of inhibition by PPi *in vivo* has so far not been addressed. Cytosolic PPi concentrations of 0.2–0.3 mM as reported for spinach leaves (Weiner et al., 1987) would clearly not be compatible with E1 activity suggesting that PPi levels are maintained at substantially lower levels at least in the immediate environment of E1 enzymes. Many nuclear proteins are modified by SUMOylation (Rytz et al., 2018) and the SUMO conjugation complex self-assembles into nuclear bodies (Mazur et al., 2018). Information regarding the nuclear concentration of PPi is lacking, but the fact that DNA and RNA synthesis occurs against such high concentrations of PPi argues not only that soluble pyrophosphatases play an important role in the nucleus but could also suggest that nuclear PPase activity is higher than in the cytosol. However, the fact that a quadruple knockout mutant lacking four of five PPa-isoforms showed no obvious phenotype whereas the combined loss of the H^+^-PPase AVP1 and a single PPa-isoform causes severe dwarfism due to high PPi concentrations (Segami et al., 2018) shows clearly that cytosolic and nuclear pools of PPi are controlled by the combined action of soluble and H^+^-PPase.

At least for plants, converting the substantial energy present in PPi into a proton-gradient seems preferable to releasing it as heat and the soluble PPases might thus only function as emergency valves. But is accumulation of PPi to inhibitory levels only occurring in mutant backgrounds or is there evidence that it is actively prevented under stress conditions in the wild-type? The fact that four PPa-isoforms are transcriptionally up-regulated during the first six hours of the cold acclimation response (Figure 2) indicates that control of PPi levels is an integral part of the cold stress response and it remains to be determined if this is also true for other responses in particular for heat stress. Constitutive overexpression of AVP1 has been shown to cause increased growth of diverse crop plants under various abiotic stress conditions. Greater vacuolar ion sequestration, increased auxin transport, enhanced heterotrophic growth, and increased source to sink transport of sucrose have all been proposed to explain individual aspects of the phenotypes observed in plants lacking or over-expressing AVP1 (Park et al., 2005; Pasapula et al., 2011; Schilling et al., 2017; Yang et al., 2014). Here, we propose modulation of SUMOylation by cellular pyrophosphate levels as a unifying hypothesis that might explain both, the stress-related as well as the developmental aspects of the multifaceted AVP1 loss- and gain-of function phenotypes. Although our hypothesis needs further experimental validation in particular regarding the developmental phenotypes, it seems obvious that a combination of tissue-specific and inducible expression of PPi-hydrolysing enzymes might turn out to be an efficient way of generating stress-tolerant crops for the future.

## EXPERIMENTAL PROCEDURES

### Plant material and growth conditions

*Arabidopsis thaliana* Col-0 ecotype was used in all experiments as control. The two V-PPase mutant lines *fugu5–1* and *fugu5–3* and the yeast soluble pyrophosphatase overexpression line (*AVP1:IPP1 / fugu5–1*) were previously described (Ferjani et al., 2011). Transgenic *UBQ:AVP1 #18–4* was described in (Kriegel et al., 2015). *ice1–2* (SALK_003155) mutant was obtained from the SALK population (http://signal.salk.edu; Alonso et al., 2003). Seeds were surface sterilized with ethanol and stratified for 48h at 4°C. For the heat shock tolerance assays, seedlings were grown on plates with standard growth medium (0.5% Murashige and Skoog (MS), 0.5% phyto agar, and 10mM MES, pH 5.8) for 10 days under long day conditions (16 h light/8 h dark) at 22°C at 125 μmol·m^−2^·s^−1^. At day 10, treatment plates were exposed to 40°C for 4 hours while the control plates were kept in growth conditions. For freezing tolerance assay, electrolyte leakage assay, PPi and sugar determination and real time RT-PCR, plants were grown for 6-weeks on soil under short day conditions at 22°C at 125 μmol·m^−2^·s^−1^ Afterwards they were cold acclimated for 4 days at 4°C. Untreated plants were maintained in the same conditions as the growth period. To determine SUMO and ICE1 protein amounts upon cold and heat treatments, seedlings were grown in liquid culture (0.5% Murashige and Skoog (MS), 0.5% sucrose, 10mM MES, pH5.8) under long day conditions (16 h light/8 h dark) at 22°C at 125 μmol·m^−2^·s^−1^. Growth period was 10 days in 50ml liquid culture in a 300ml flask on a horizontal shaker with 100 rpm speed. After 10 days, part of flasks was either subjected to 30 min 40°C or 3 hours 4°C. Control samples were kept at growth conditions.

### Construct design and plant transformation

*UBQ:PPa5-GFP* construct was generated using GreenGate (GG) cloning (Lampropoulos et al., 2013). The 1097 base pairs coding sequence of *PPa5* was amplified from Arabidopsis thaliana Col-0 cDNA with primers listed in table 1, attaching BsaI recognition sites and specific GG-overhangs. To prevent cutting, the internal BsaI site was mutated by site directed mutagenesis. Thereafter, the PCR product and the empty entry module (pGGC000) were digested with BsaI, the digestion was purified and then ligated. After test digestion positive clones were checked by sequencing. The final construct was assembled in a GG reaction from modules listed in table 2 and transformed into Agrobacterium tumefaciens ASE strain harbouring the pSOUP plasmid. Arabidopsis thaliana ecotype Col-0 and *fugu5–1* plants were used for transformation via floral dipping (Clough and Bent, 1998).

**Table 1.**
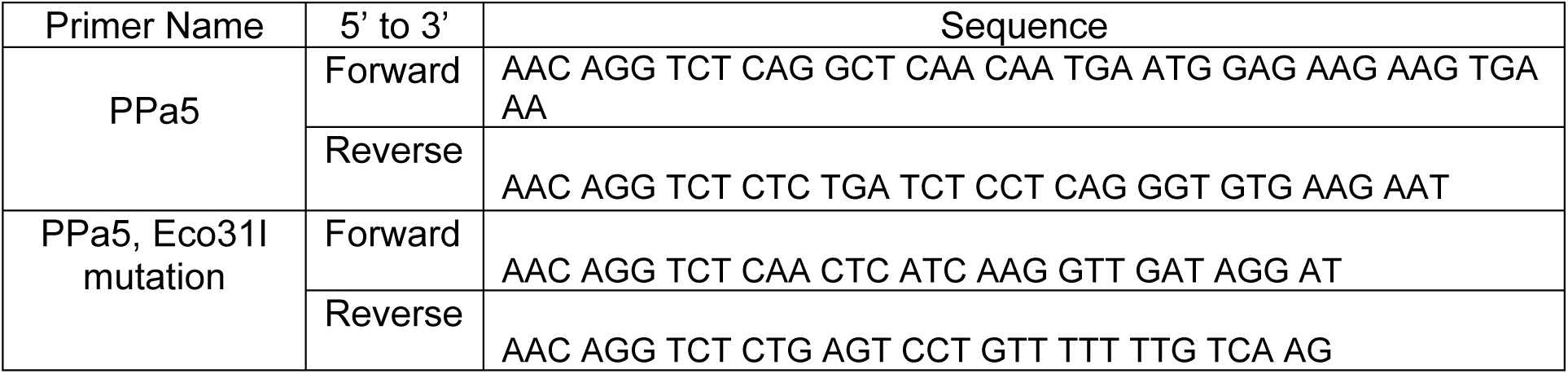
Primers of *UBQ:PPa5-GFP* construct

**Table 2.** List of GG modules

| GG module name | Type | AGI | Reference |
| --- | --- | --- | --- |
| pGGA006 | *UBIQUITIN 10* (*UBQ10*) promoter | AT4G05320 | Lampropoulos et al., 2013 |
| pGGB003 | B-dummy | na | Lampropoulos et al., 2013 |
| pGGC-PPa5 | *PPa5* | AT4G01480 | This work |
| pGGD001 | Linker-GFP | na | Lampropoulos et al., 2013 |
| pGGE001 | *RBCS* terminator (from pea) | na | Lampropoulos et al., 2013 |
| pGGF012 | *pUBQ10:Hyg<sup>R</sup>:tOCS* | na | Lampropoulos et al., 2013 |
| pGGZ001 | Vector backbone | na | Lampropoulos et al., 2013 |

### Yeast strain generation and growth

To replace the endogenous IPP1 promoter with the inducible GAL1 promoter and simultaneously introduce an N-terminal HA-tag, plasmid pFA6a-His3MX6-PGAL1 was amplified with primers Ipp1-F4/Ipp1-R3 (Longtine et al., 1998). The resulting PCR product was used for transformation of a wild type S. cerevisiae W303 strain (SSY122; (Szoradi et al., 2018). Correct promoter replacement in the resulting IPP1prΔ::HIS3-GAL1pr-HA-IPP1 strain (SSY2542) was confirmed by colony PCR and lack of growth on glucose-containing medium. Cells were grown on synthetic complete medium (CSM –Uracil (MP #4511212), Difco™ yeast nitrogen base (BD #233520), Uracil (Sigma #U1128), Adenine Hemisulfate (Sigma #A9126)) supplemented with appropriate carbon sources. All determinations were done on exponentially growing cells (A_600_ ≤ 0.5). To maintain cultures for several hours below an A_600_ of 0.5, they were diluted with fresh medium every two hours until the end of the experiment (semi-continuous culture). A pre-culture containing galactose was grown at 28°C shaking until A_600_ 0.5, then divided to four, for temperature and carbon resource manipulation: Glucose / 28°C, Galactose / 28°C, Galactose / 40°C, Glucose / 40°C. Samples were taken at indicated time points (0, 6, 15 hours). For heat treatment samples were taken from 28°C one hour before the indicated time point and incubated at 40°C for an hour.

### Protein preparation and immunoblotting analysis

To determine total sumoylation amount in Arabidopsis, total proteins were extracted from liquid grown wt, V-PPase mutants, *UBQ:PPa5/fugu5–1* and *UBQ:AVP1* as described in (Castaño-Miquel et al., 2013). 15 μg protein was loaded to 7.5%, 1.5 mM SDS-gels. Anti-SUMO1 (1:1000; Agrisera) was used as primary antibody. To measure ICE1 protein, liquid grown Col-0 and *fugu5–1* and *ice1–2* were used for total protein extraction as described in Castano-Miquel et al., 2013. Anti-ICE1 (1:1000; Agrisera) was used as primary antibody. Anti-cFBPase (Agrisera; 1:5000) was used as loading control for both SUMO and ICE1. For all immunoblots, HRP-anti-rabbit was used as secondary antibody (1:10000; Promega). To determine the amount of soluble pyrophosphatase and the amount of total sumoylated proteins in yeast, total protein extraction was performed as described in Szoradi et al., 2018 with addition of 20 mM NEM. 15 μg protein was loaded to 7.5%, 1.5 mM SDS-gels. Anti-IPP1 (ABIN459215, Antibodies-online GmbH, 1:1000) and anti-SUMO1 (1:1000; Agrisera) were used as primary antibodies. HRP anti-rabbit was used as secondary antibody. Anti-PGK1 (Abcam, 1:100000) was used as the loading control. An anti-mouse antibody was used as secondary antibody (GE Healtcare UK, 1:5000).

### Determination of PPi and soluble sugar levels via ion-chromatography

6-weeks old short day grown rosettes were ground in liquid nitrogen and aliquots of ~200-400 mg were used to quantify PPi and soluble sugars. Compounds were extracted with 0.5 ml ultra-pure water for 20 min at 95°C with vigorous shaking, and insoluble material was removed by centrifugation at 20,800 g for 20 min. PPi was measured using an IonPac AS11-HC (2 mm, Thermo Scientific) column connected to an ICS-5000 system (Thermo Scientific) and quantified by conductivity detection after cation suppression (ASRS-300 2 mm, suppressor current 29–78 mA). Prior separation, the column was heated to 30°C and equilibrated with 5 column volumes of ultra-pure water at a flow rate of 0.3 ml/min. Soluble sugars were separated on a CarboPac PA1 column (Thermo Scientific) connected to the ICS-5000 system and quantified by pulsed amperometric detection (HPAEC-PAD). Column temperature was kept constant at 25°C and equilibrated with five column volumes of ultra-pure water at a flow rate of 1 ml min-1. Data acquisition and quantification was performed with Chromeleon 7 (Thermo Scientific).

### Freezing tolerance assay

Plant freezing tolerance was determined with 6-weeks old short-day grown plants. For cold acclimation, 6-week-old plants were incubated at 4°C for 4 days with same photoperiod. Non-acclimated plants were kept at 22°C during this period. Plants were wetted thoroughly to promote freezing, then placed in a controlled temperature chamber (Polyklima, MN2-WLED). First they were kept at 0°C for 1h. Afterwards, they were subjected to temperatures from −1 to −10°C, reduced 1°C every 30 min. After thawing at 4°C overnight, plants were moved back to 22°C. Images were taken before cold treatment and 1 week after the freezing treatment. Dead and alive leaves were counted after the photos were taken.

### Electrolyte leakage from leaves

Electrolyte leakage was measured from fully developed rosette leaves of 6-week-old plants. For each temperature five leaves were collected from each genotype. Each leaf (5th or 6th rosette leaf) was placed into a tube containing 3 mL deionized water, then placed to 0 °C at a temperature-controlled climate chamber. Temperature was decreased by 2 °C every hour. At −2 °C an ice chip was added to initiate nucleation. Tubes were collected at −2, −4, −6, −8 and −10 °C and placed to 4 °C to thaw overnight. Next day 2 ml deionized water was added and tubes were placed overnight on a horizontal shaker (100 rpm) at 4 °C. Conductivity after freezing was measured with a conductivity meter (Mettler-Toledo, FiveEasy), which was calibrated with the Mettler-Toledo Buffer solution 1413 μS. Then, samples were placed to a 100 °C water bath and boiled for 2 hours. Conductivity was again measured after boiling. Ion leakage was determined as the percent ratio of the measurement of conductivity before and after boiling.

### RNA isolation and cDNA synthesis

For the analysis of transcript levels 6 weeks old Col-0, *fugu5–1* and *UBQ:PPa5-GFP/fugu5–1* was collected after exposure to 4°C for indicated time points. RNA was isolated using the RNeasy Plant Mini Kit (Qiagen) according to manufacturer’s instructions. cDNA was synthesized from 1 μg of total RNA using M-MuLV reverse transcriptase (Thermo) and an oligo dT primer.

### Real-time RT PCR

For quantitative analysis of gene expression real-time RT PCR was applied. cDNA samples were diluted 1:50 in nuclease-free water. Real-time PCR reactions were performed using the DNA Engine Opticon System (DNA Engine cycler and Chromo4 detector, BioRad) and SG qPCR mastermix 2X (Roboklon). The real-time PCR reaction mixture with a final volume of 20 μl contained 0.5 of each forward and reverse primer, 10 μl SYBR Green Mix, 4 μl cDNA and 4 μl of RNase-free water. The thermal cycling conditions were composed of an initial denaturation step at 95°C for 15 min followed by 40 cycles at 95°C for 15 sec, 60°C for 30 sec and 72°C for 15 sec and ended with a melting curve. For the analysis of each sample three analytical replicas were used. Target genes were normalized to the expression of *Actin2*. Primer sequences are listed in Table 2.

**Table 3.**
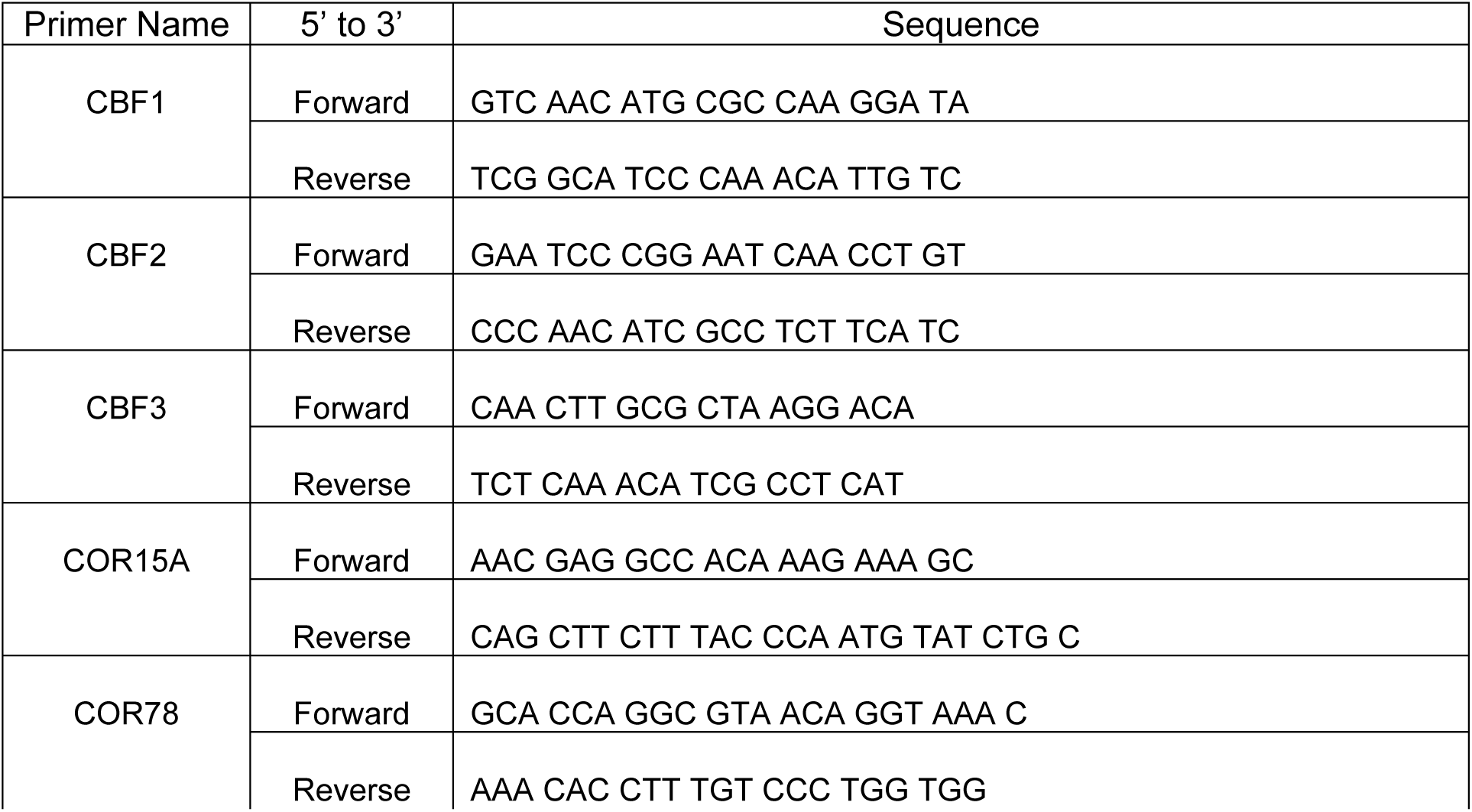
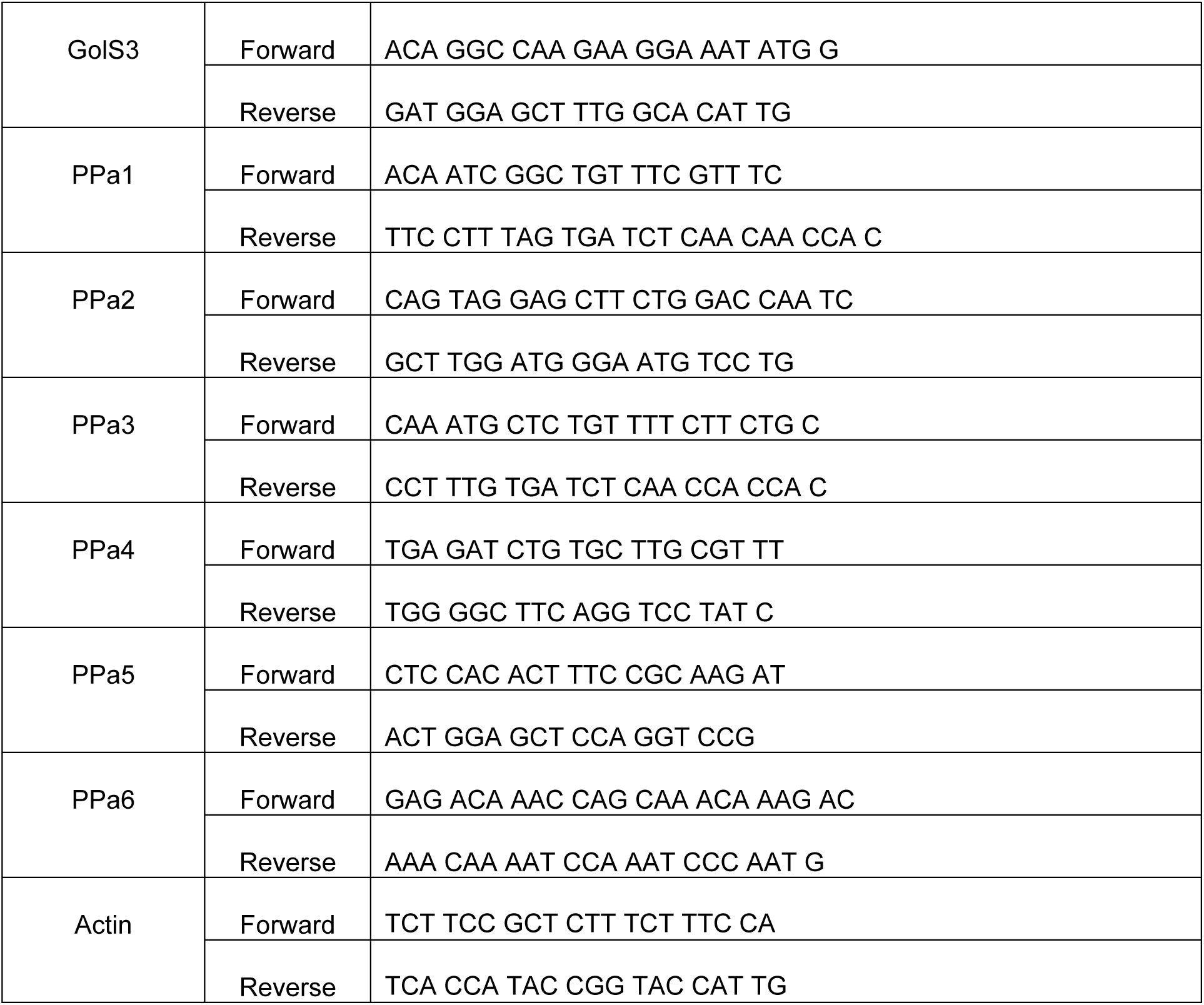
qRT primers

**Table 3.**
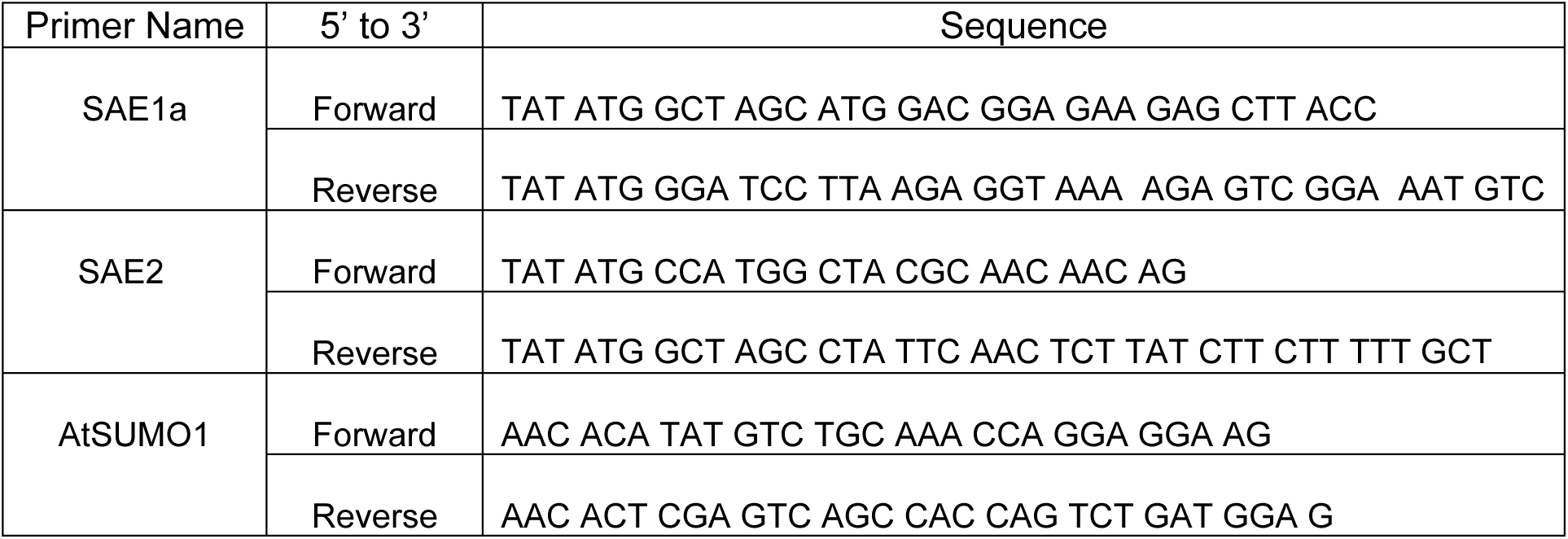
Primers for protein purification

### In vitro FRET-based sumoylation assay

Sumoylation of CFP-RanGAPtail with YFP-SUMO was carried out using a FRET-based high-throughput assay as previously described with minor changes (Werner et al., 2009)Bossis et al., 2005). Final concentrations of the FRET components were Uba2/Aos1 (E1, 20 nM), Ubc9 (E2, 30 nM), YFP-SUMO3 and CFP-RanGAPtail (300 nM). ATP substrate was prepared as a stock solution of 300 mM ATP-BTP (pH 8.0). 1 mM ATP was used for the assays unless stated otherwise. For the PPi application, 30 mM PPi-BTP (pH 7.5) stock solution was prepared. To determine the effects of PPi hydrolysis on the FRET assay, 0.8U E. coli inorganic pyrophosphatase (NEB) was used and the buffer solution that the pyrophophatase includes was added to the control wells (20 mM Tris-HCl, 100 mM NaCl, 1 mM Dithiothreitol, 0.1 mM EDTA, 50% Glycerol, pH 8.0). Michealis-Menten fittings and Vmax and Km calculations were done in Origin software according to the ATP titration (0–10 μM) performed with different PPi concentrations (0, 7.5 and 15 μM).

### Cloning, expression and protein purification of Arabidopsis E1 ligase and SUMO1

Conjugation-competent AtSUMO1 (1–93) was amplified from cDNA, using the primers AtSUMO1-NdeI-Fw and AtSUMO1-XhoI-Rv, and cloned in the bacteria expression vector pET28b(+). The coding sequence of SAE1a was amplified from A. thaliana cDNA ligated into pET28a via NheI and BamHI sites in-frame behind the coding sequence for a 6xHis-tag. SAE2 was amplified from A. thaliana cDNA and ligated into the pCR™-Blunt II-TOPO^®^ vector. pET11d was cut with BamHI and the TOPO-vector was cut with NheI. Both linear DNA fragments were blunt ended with T4 polymerase. Both DNA fragments were subsequently restricted with NcoI. The DNA fragment carrying the SAE2 coding sequence was ligated into pET11d via the NcoI cohesive end and the blunt end. Recombinant proteins were purified as previously described (Werner et al., 2009).

### *In vitro* E1-Thioester Assay

E1-Thioester assay was performed as previously described in Castaño-Miquel et al., 2013 with addition of final concentrations of 1–10 μM PPi. 1 mM of ATP was used for all reactions unless stated otherwise.

## ACKNOWLEDGMENTS

We are grateful to Beate Schöfer, Barbara Jesenofsky and Anne Newrly for excellent technical assistance. This work was supported by DFG grants SCHU1151/13–1 to KS, AN1323/1–1 to ZA and SFB1036 (TP15) to FM. SS acknowledges support through DFG grant EXC 81.

---

**Supp. Figure 1.**
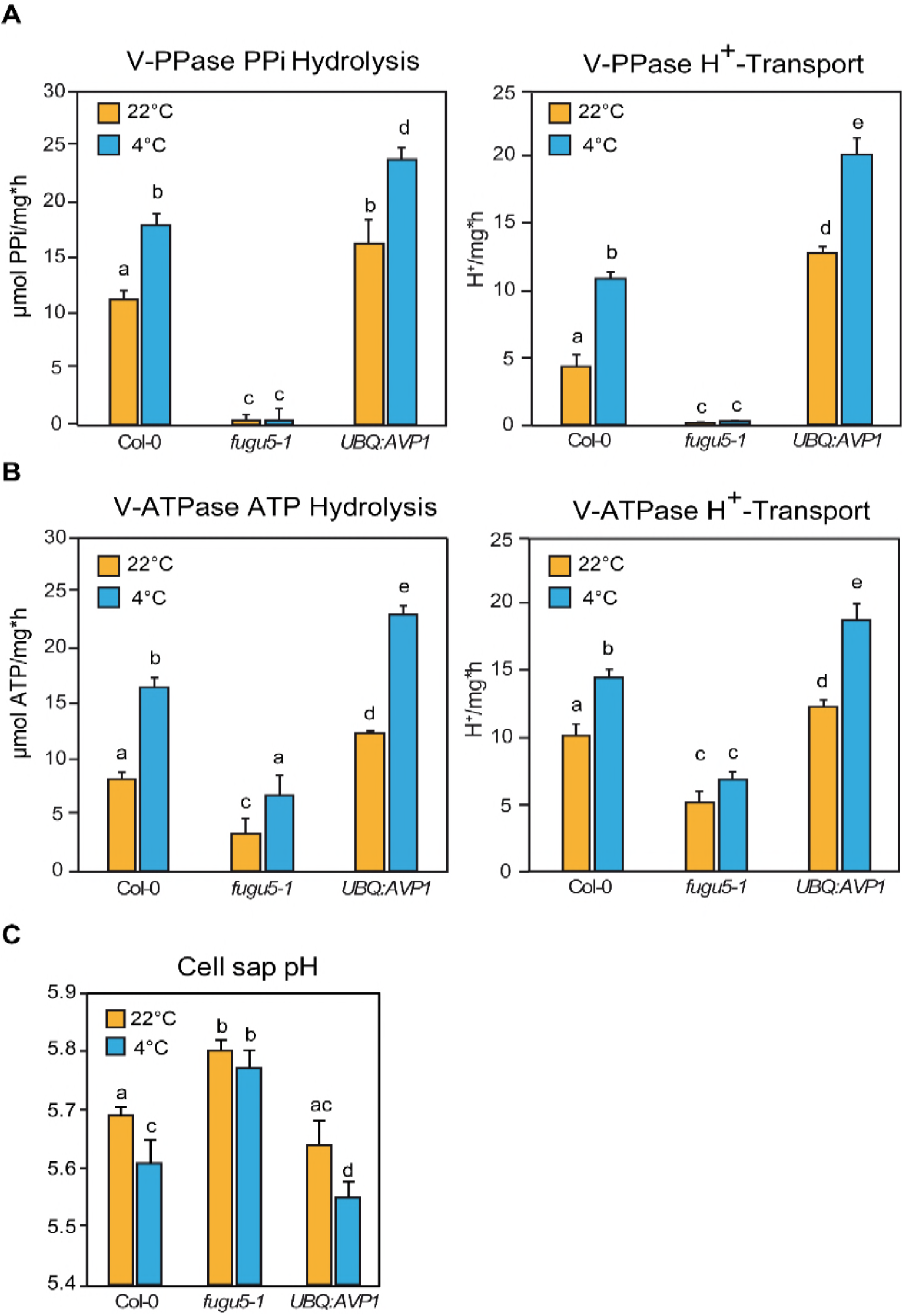
Increased Proton Transport Activity of V-ATPase upon Cold Acclimation Requires an Active V-PPase. **(A)** Enriched tonoplast proteins were used to determine K^+^-stimulated PP_i_ hydrolysis and H^+^ transport activity of V-PPase. **(B)** Enriched tonoplast proteins were used to determine KNO_3_-inhibited ATP hydrolysis and H^+^ transport activity of V-ATPase. **(C)** Cell sap pH measurement of rosette leaves. Error bars show SD of the mean with n=12 samples of 3 biological replicates. **(A-C)** Col-0, *fugu5–1* and *UBQ:AVP1* were grown for 6 weeks under short-day conditions then were cold acclimated for 4 days at 4°C. Untreated plants were maintained in the same conditions as the growth period. Error bars represent SD of n=3 biological replicates. Significant differences are indicated by different letters (Two-way ANOVA followed by Tukey’s test, p<0.05).

**Supp. Figure 2.**
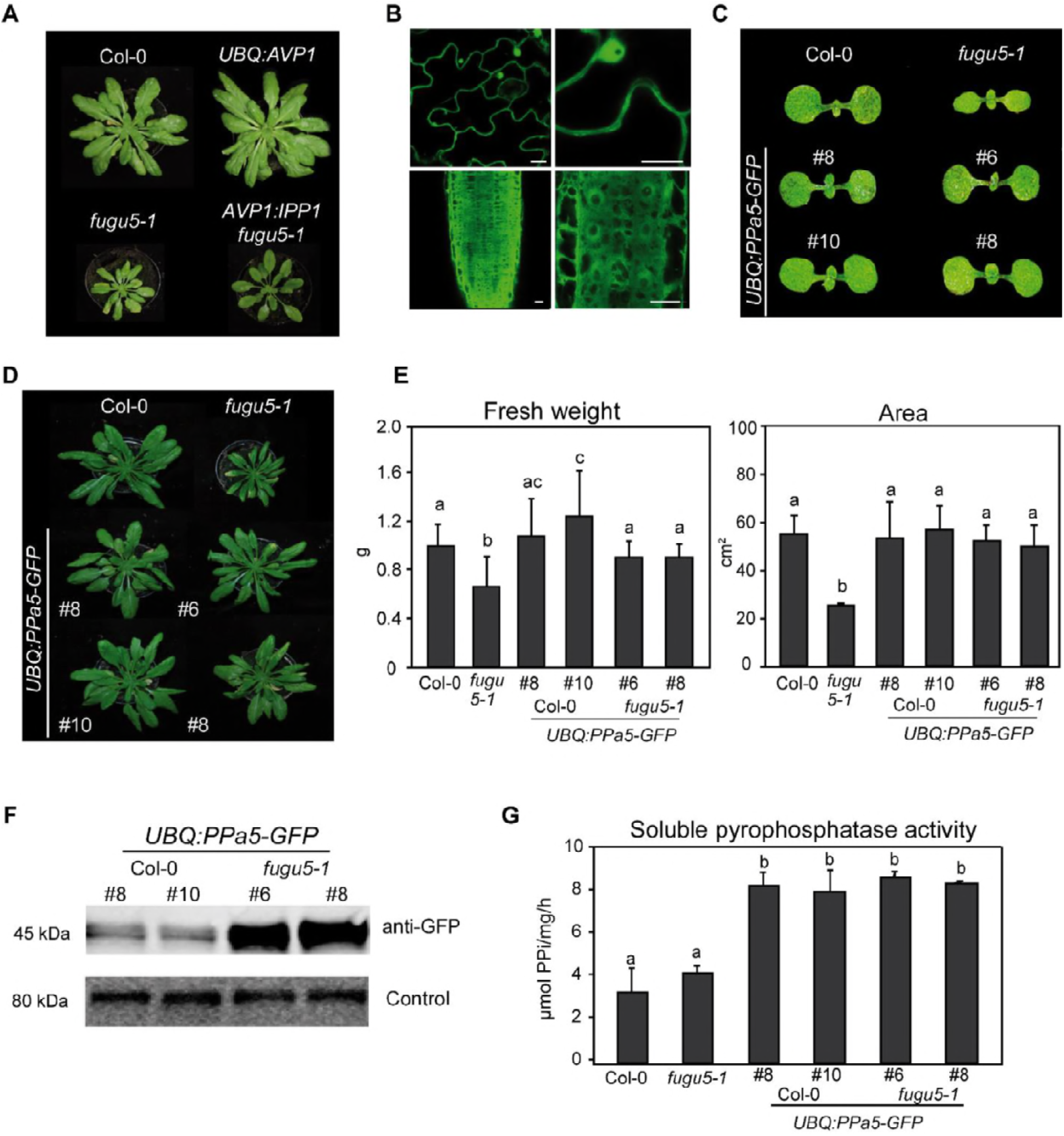
Overexpression of the Arabidopsis soluble pyrophosphatase PPa5 is sufficient to complement *fugu5–1*. **(A)** Comparison of 6-week old rosettes phenotypes of Col-0, *fugu5–1* and overexpression lines of Arabidopsis vacuolar pyrophosphatase AVP1 and yeast soluble pyrophosphatase IPP1. **(B)** Representative images showing the localization of *UBQ:PPa5-GFP* to cytosol and nuclues in shoot (upper panel) and root cells (lower panel). Scale bars: 10μm. **(C)** Cotyledon phenotypes of 5 days old seedlings of Col-0, *fugu5–1* and *UBQ:PPa5-GFP* in Col-0 and *fugu5–1* backgrounds. **(D)** Comparison of 6-week old short day grown rosette phenotypes of Col-0, *fugu5–1* and *UBQ:PPa5-GFP* in Col-0 and *fugu5–1* backgrounds. **(E)** Measurements of fresh weight and whole rosette area of Col-0, *fugu5–1* and *UBQ:PPa5-GFP* in Col-0 and *fugu5–1* backgrounds. Plants were grown for 6 weeks under short-day conditions. To determine rosette area Rosette Tracker plug-in of ImageJ is used. Error bars represent SD of the mean of n=20 of 3 biological replicates. **(F)** Analysis of *UBQ:PPa5-GFP* protein amount in Col-0 and *fugu5–1* background with anti-GFP. Soluble proteins from 6-week old rosettes grown under short-day conditions were extracted. An internal control provided by SPL detection kit (DyeAGNOSTICS) was used for normalization. One representative image from three biological replicates is depicted. **(G)** Soluble proteins of Col-0, *fugu5–1* and *UBQ:PPa5-GFP* in Col-0 and *fugu5–1* backgrounds were used to determine K^+^-stimulated PP_i_ hydrolysis. Plants were grown for 6 weeks under short-day conditions. Error bars represent SD of n=3 biological replicates. Significant differences are indicated by different letters (Two-way ANOVA followed by Tukey’s test, p<0.05).

**Supp. Figure 3.**
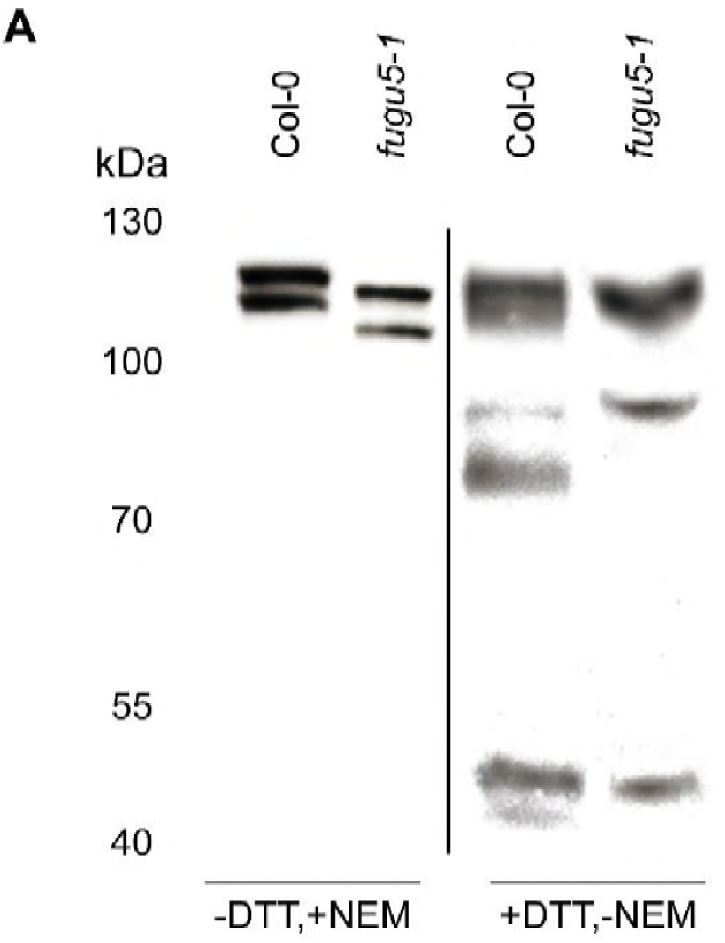
Western blot detection of ICE1 in protein extracted in the presence of DTT. **(A)** Comparison of total protein of 10 days old Col-0 and *fugu5–1* seedlings extracted +/− DTT (5 mM) and NEM (20 mM).

**Supp. Figure 4.**
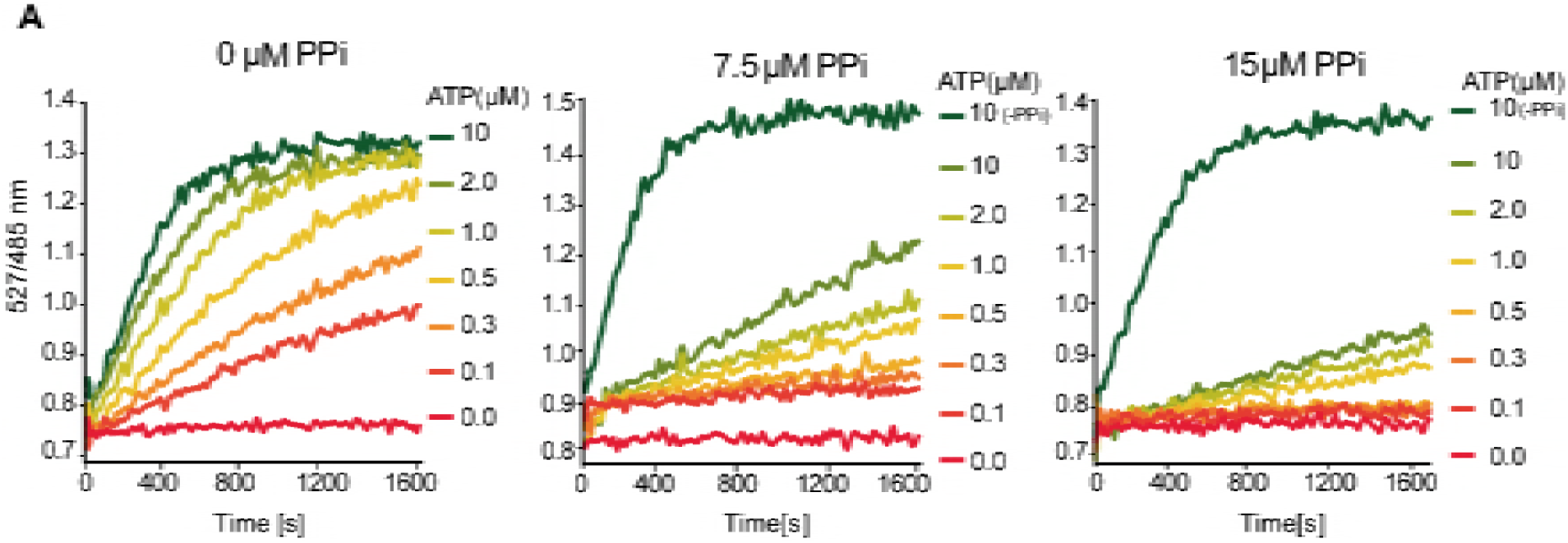
Elevated PPi concentrations leads to mixed inhibition of SUMOylation *in vitro*. **(A)** Sumoylation assays demonstrating the effect of increasing amounts of PP_i_ to the speed of the reaction in a range of 0–10μM ATP concentration. Experiments were repeated 4 times.

**Supp. Figure 5.**
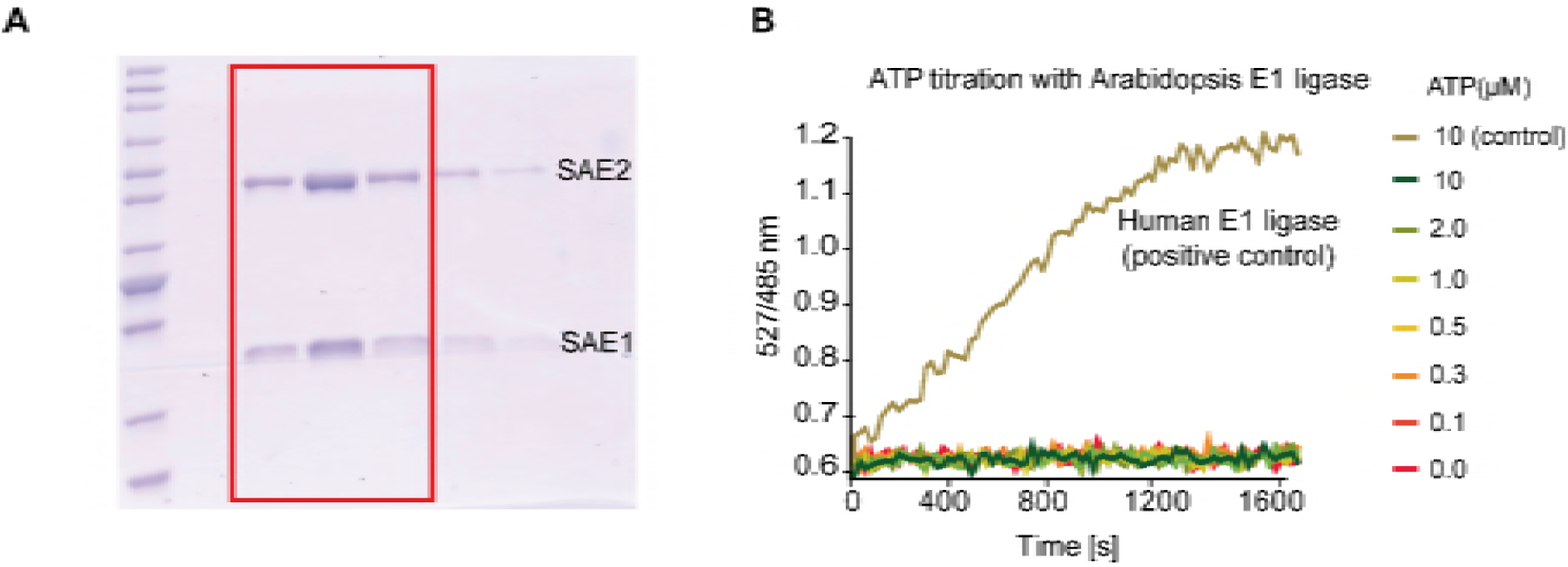
AtSAE1/2 purification and its activity in in vitro FRET based SUMOylation assay. **(A)** Gel picture after purification of AtSAE1 and AtSAE2. Highlighted fractions from the final gel filtration step were combined, dialysed and used for subsequent experiments. **(B)** The purified Arabidopsis E1 activating enzyme is not functional in the FRET based sumoylation assay. Assays were set up as described for Figure 6, but with recombinant Arabidopsis E1 enzyme. Human E1 activating enzyme was used in a positive control.

## Supplementary Material and Methods

### Plant material and growth conditions

Same plant material described in the main section was used. Seedling growth medium and conditions for the confocal imaging was the same as the conditions described for the heat shock tolerance assay. The plant material used for enzyme assays, cell sap measurements and phenotypic assays were grown the same as described for material for freezing tolerance assay. Cold acclimation conditions were also the same. For analysis of cotyledon phenotypes seeds were surface sterilized with ethanol and stratified for 48h at 4°C in 0.5% agar solution then planted on single pots individually (n=10). Pictures are taken at day 5. Liquid culture conditions to grow the material for ICE1 determination is the same as described in the main section.

### Tonoplast vesicle preparation and enzyme assays

Rosette leaf material (75 g) from plants grown under short day conditions was harvested. The leaf material was homogenized in homogenization buffer containing 0.4 M mannitol, 0.1 M Tris, 10% (vol/vol) glycerol, 3 mM Na_2_EDTA, 0.5% (wt/vol) BSA, 5% (vol/vol) PVP-10, 0.5 mM butylated hydroxytoluene, 0.3 mM dibucaine, 5 mM magnesium sulphate, 1 mM PMSF (phenylmethylsulphonylfluoride), 1.3 mM benzamidine and 25 mM potassium metabisulfite. The homogenate was filtered through two layers of miracloth and centrifuged at 10,000 g for 20 min at 4°C. The supernatant was then centrifuged at 100,000 g for 45 min at 4°C. The microsomal membrane pellet was resuspended in resuspension buffer containing 0.4 M mannitol, 6 mM Tris-MES (pH 8) and 10 (vol/vol) glycerol. Soluble part was kept for measuring the soluble pyrophosphatase activity and quantification of PPa5-GFP protein levels. Tonoplast vesicles were obtained by performing a sucrose gradient with 22% sucrose. Centrifugation was performed at 97,000 g for 2 hours. Protein concentrations were determined as reported previously (Bradford, 1976). ATP and PP_i_ hydrolysis was measured at 28°C as described previously (Krebs et al., 2010). Same method was also used for measuring soluble pyrophosphatase activity with soluble proteins. The ATP and PP_i_-dependent proton transport activities were estimated from the initial rate of ATP-dependent fluorescence quenching in the presence of 3 mM ATP using the fluorescence dye ACMA (9-Amino-6-Chloro-2-Methoxyacridine) with 50μg enriched tonoplast protein. Excitation wavelength was 415 nm, and emission was measured at 485 nm in Jasco fluorescence spectrometer. V-PPase H^+^ transport medium includes 25mM HEPES-BTP (pH 7.2), 250 mM Sorbitol, 1.5 mM MgSO_4_, 50 mM KCl and 0.3mM PP_i_-BTP (pH 7.5) final concentration in 1 ml volume. V-ATPase H^+^ transport medium includes 10 mM ATP-MES (pH 8.0),0.25 M Mannitol, 3 mM MgSO_4_, 100 mM TMA-Cl and 1.5 mM ATP-BTP (pH 7.5) final concentration in 1 ml volume.

### SDS-PAGE and immunoblotting analysis

To determine protein levels of V-ATPase and V-PPase at 22 and 4°C in tonoplast membrane extracts of Col-0, *fugu5–1* and *UBQ:AVP1*. The primary antibody against the V-PPase was purchased from Cosmo Bio (1:10,000) and the primary antibody against VHA-C is as previously described (Schumacher et al., 1999). To determine the *UBQ:PPa5-GFP* levels in Col-0 and *fugu5–1* background soluble proteins were extracted as described in tonoplast vesicle preparation section. Anti-GFP (Agrisera, 1:10000) was used as primary antibody. An internal control from SPL kit was used for normalization (NH, DyeAgnostics). To measure ICE1 protein, Col-0 and *fugu5–1* were grown in liquid culture. Material was split in two for total protein extraction, same buffer described in Castaño-Miquel et al., 2013 was used for one part, and same buffer without NEM and with addition of 5 mM DTT used for the other. Anti-ICE1 (1:1000; Agrisera) was used as primary antibody. For all immunoblots, HRP-anti-rabbit was used as secondary antibody (1:10000; Promega). Imaging was carried out using a cooled CCD camera system (Intas ADVANCED Fluoreszenz u. ECL Imager). Western blots were quantified with Fiji (based on ImageJ 1.47t).

### Cell sap pH measurements

Cell sap pH measurements were conducted as previously described (Krebs et al., 2010)

### Confocal Microscopy

Localization of *UBQ:PPa5-GFP* construct was determined using a Leica TCS SP5II microscope equipped with a Leica HCX PL APO lambda blue 63.0 3 1.20 UV water immersion objective. GFP was excited at 488 nm using a VIS-argon laser. Fluorescence emission of GFP was detected between 500 and 555 nm. The Leica Application Suite Advanced Fluorescence software was used for image acquisition. Post processing of images were performed using Fiji.

